# Resolving the paradox of local warning signal diversity with predator learning

**DOI:** 10.1101/2023.05.04.539348

**Authors:** Chi-Yun Kuo

## Abstract

Coexistence of distinct warning signals at local scales has long stood as a paradox, as selection is expected to preserve only the most common signal. So far, there has not been an explanation that is both broadly applicable and testable. This study presents a novel and generalizable resolution to this paradox (the Unforgetful Predator Hypothesis) by showing that prey displaying a rarer warning signal can persist if predators have low enough forgetting rates relative to prey generation time. In addition, inducing a high level of predator avoidance facilitates warning signal diversity when prey do not compete. In the presence of prey competition, however, warning signal diversity is more likely to occur if prey elicit intermediate levels of avoidance, such that the competitive disadvantage for individuals displaying the rare signal can be offset by predation. This hypothesis can be tested by quantifying predator avoidance and forgetting rate in laboratory and field experiments. As the level of predator avoidance is also crucial in determining the fate of rare signal in communities, I performed a meta-analysis to examine the determinants of its variation and found that higher predator avoidance can be observed in the following situations – when prey unprofitability is due to toxicity rather than mere unpalatability, when predators search widely for prey and/or prey aggregate, and when predators could differentiate between unprofitable and profitable prey using only color or pattern. In addition to resolving the paradox, these findings help inform the types of communities in which distinct warning signals can stably coexist.

**Significance:** Coexistence of distinct warning signals within a community represents an evolutionary paradox that still awaits a resolution that is both general and testable. I used ecologically realistic simulations to show that warning signal diversity can occur if predators have long enough memory and if prey elicit either low or moderate levels of avoidance, depending on whether they compete. A meta-analysis further shows that higher levels of predator avoidance tend to occur when unprofitable prey are highly unprofitable, when predators search widely for prey, when prey aggregate, and when prey profitability could be discerned by only color or pattern. These findings offer a testable resolution to the paradox and inform the types of communities where warning signal diversity may occur.

## Introduction

Unprofitable prey often evolve conspicuous colorations that advertise their defenses (i.e., aposematism, (1). These visual advertisements act as honest warning signals that predators can use to avoid these prey through associative learning. The widespread occurrence of warning signals has inspired numerous hypotheses regarding their evolution and maintenance in nature (2–4). Among these, Müller’s pioneering idea and subsequent theoretical developments aptly explains the often-observed signal convergence among unprofitable prey (i.e., Müllerian mimicry) as the result of positive frequency-dependent selection (hereafter FDS). We therefore expect aposematic prey, at least those targeted by the same predators, to be homogeneous in signal appearance at local scales (ref. 5 and references therein). Yet, it is by no means rare to observe distinct warning signals coexisting stably within communities (6, 7). The simultaneous occurrence of warning signal convergence and diversity at local scales represents a stark contrast from theoretical expectations and has long stood as a paradox.

One potential resolution to this paradox is that species displaying distinct warning signals might segregate along certain microhabitat axes, and predators learn to avoid different signals in each microhabitat type (8, 9). This possibility can be tested empirically but so far seems only able to unambiguously explain warning signal diversity in Costa Rican ithomiine butterflies, in which distinctively colored species fly at different height strata due to host plant distribution (10, 11). Notably, microhabitat segregation does not appear to be clearcut between distinctly colored *Heliconius* butterflies or *Oophaga* poison frogs (12, 13), indicating that other mechanisms may be responsible. There have also been hypotheses seeking to directly explain local warning signal diversity with predator behavior and learning. A theoretical study used Müller’s original idea of a fixed number of prey mortality for predator education and found that if the prey population is large enough compared to the fixed mortality, the strength of FDS would be substantially weakened (5). Further modeling predator learning with a Pavlovian process (14–20), another study found that a large number of distinct warning signals converged into a smaller number of mimicry rings (21), likely due to weak FDS from large prey population sizes (7). These findings insightfully showed that reduced predation can be a key facilitator for warning signal diversity, but they explored this condition as a consequence of prey abundance rather than predator learning.

A significant advancement in explaining local warning signal diversity with predator learning was the use of a Bayesian process to model the development of predator avoidance. The Bayesian learning framework was formulated by (22) as an alternative to the Pavlovian learning models to better account for the adaptive nature of foraging decision-making (23). Bayesian learning describes predator avoidance as the outcome of an exploitation-exploration tradeoff, in which each interaction with defended prey with a certain appearance leads the predator to more accurately estimate their overall profitability. When predators no longer benefit from sampling more prey, a complete avoidance results (i.e., an attack probability of zero). A Bayesian predator that encounters a novel prey type that are both rare and potentially highly unprofitable are expected to not sample such prey altogether, resulting in persisted neophobia that favors the evolution of warning signal polymorphism in defended prey (22, 24). The occurrence of this persistent neophobia hinges on the ratio between the cost incurred if the attacked prey is defended and the benefit if the attacked prey is undefended, as well as the carrying capacity of the prey population (24). The Bayesian learning framework offered a theoretical breakthrough for explaining local warning signal diversity with predator learning, but it has two limitations. First, the Bayesian learning framework does not consider predator forgetting. More importantly, it is difficult, if not impossible, to test whether warning signal diversity is maintained by Bayesian predators, as the abovementioned costs and benefits of prey consumption can be hard to quantify. Furthermore, in Müllerian systems where all prey are defended, the benefit of consuming an undefended prey might be neither biologically meaningful nor estimatable.

Here, I present a novel explanation for local warning signal diversity that directly relates to predator learning and forgetting (hereafter The Unforgetful Predator Hypothesis). This hypothesis posits that predators being sufficiently unforgetful is key to maintaining warning signal diversity for two reasons. First, forgetful predators may repeatedly impose the cost of predator education, leading to more frequent and persistent occurrences of positive FDS (25). Moreover, when predators are quick to forget learned avoidance, attack probability and the resulting prey mortality will be higher than what would be expected from prey profitability alone. These disadvantages are particularly severe for rare warning signals, as their low encounter rates lead to more opportunities for forgetting. When predators can retain learned avoidance for long enough, the mortality costs for predator education are rarely paid and thus become trivial compared to baseline predation and reproduction. Stable persistence of rare warning signals can then be achieved as a balance between reproduction and predation, with the latter being determined by dynamic changes in attack probabilities due to prey profitability, predator learning, and forgetting. I argue that even though the Pavlovian learning framework may offer limited mechanistic insights for avoidance learning, it is a more suitable phenomenological model for testing The Unforgetful Predator Hypothesis for its explicit incorporation of forgetting, and the fact that the key parameters, namely predators’ final attack probability after learning is complete and forgetting rate, can be readily quantified with standardized experiments. Even though a density-independent final attack probability from predators has been criticized (e.g., (3, 5, 26), the proportions of prey attacked by predators do appear to approach an asymptote with repeated interactions in learning experiments (see references in *Meta-analysis of the variation in final attack probability*).

Using the Pavlovian learning framework, I also considered the effects of two important phenomena of predator learning – cognitive generalization and social learning – in facilitating local warning diversity. Cognitive generalization can occur when a novel warning signal resembles a learned signal (e.g., refs. 27–29). Generalization could have two important effects. First, predators may avoid a novel signal to some extent without prior exposure. Second, experiences with each signal types are no longer independent, such that interactions with one signal can cause a “carry-over” effect on attack probabilities towards the other signal. Social learning describes the phenomenon that predators can learn from observing the negative responses of other con- or heterospecific individuals towards prey (30–32). When predators can learn socially, experiences of individual predators are not independent, and avoidance can develop more quickly in predator populations. Generalization and social learning are also important because direct encounters with unprofitable prey is no longer necessary to reinforce learned avoidance and prevent forgetting. These two phenomena have been considered in studies that modeled predator learning (21, 33–35), yet their facilitative effects on local warning signal diversity have never been explicitly tested. I constructed both deterministic population-level models and the individual-based models to elucidate the relationship between predator learning and warning signal diversity, while accounting for generalization and social learning,

Since the final attack probability is a key parameter of the Pavlovian learning framework and directly determines the baseline predation even after learning is complete, I performed a meta-analysis to examine the variation in this value and the implications on the likelihood of local warning signal diversity in natural communities. For this, I surveyed the avoidance learning literature, focusing on the roles of the following predictors: the reason for prey unprofitability, the manner in which predators encountered prey, and the features that distinguished profitable and unprofitable prey. Specifically, I tested the following hypotheses and predictions. First, the degree of prey unprofitability determines predator avoidance; predators would have lower attack probabilities towards more unprofitable prey, such as those that are toxic or highly bitter. Second, the manner of prey encounter would affect the knowledge of prey distribution and availability and consequently predators’ attack probability. When predators encounter profitable and unprofitable prey simultaneously, predators would attack unprofitable prey less due to the knowledge that suitable prey are also present. Third, the complexity of signals or cues differentiating prey types would affect their efficacy in predator education and therefore prey avoidance. Other things being equal, signals or cues that are more effective in educating predator would result in lower attack probabilities. One could argue that multi-component features might be more effective for predator education, as these offer more precise information to help predators differentiate between prey types. If this is true, more complex signals would result in lower final attack probabilities. Yet, this assumes that predator have the cognitive capacity for using complex information. If this is not true, more complex features would confuse predators, resulting in lower educational efficacy and higher final attack probabilities.

## Results

### Unforgetful predators can maintain local warning signal diversity, especially with generalization and social learning

In the population-level model, higher initial abundance, higher intrinsic rate of increase, and low predator efficiency made the circumstances more favorable for the rare signal (Fig. 1). Prey with the rare signal were also more likely to persist if they were allowed to reach the same abundance as the common prey (Figs. S2 and S3). Importantly, sufficiently low predator forgetting rate was the main limiting factor, whereas favorable final attack probability values depended on the competitive dynamics between prey species (Fig. 1). When prey did not compete, warning signal diversity occurred at low final attack probabilities. When prey did compete, the condition for warning signal diversity was much more restricted, even though forgetting rate was still the main limiting factor in the population-level model (Figs. 1C and 2C). With prey competition, it was moderate final attack probabilities that facilitated waring signal diversity (Fig. 1). This resulted because under these attack probabilities predation sufficiently offset the competitive advantage of the common prey, thereby allowing both prey types to coexist, albeit in low numbers (Fig. S4).

**Figure 1.**
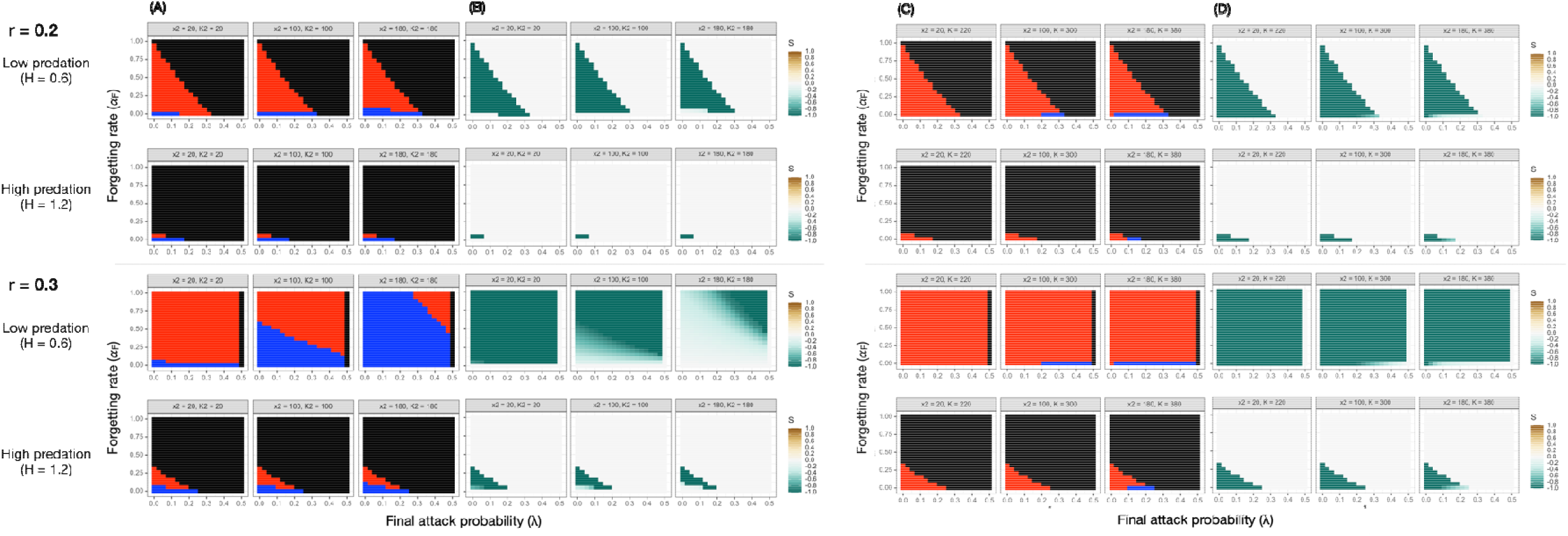
Prey persistence (A, C) and corresponding selection (B, D) on the rare prey, with respect to final attack probability (λ, x-axis) and forgetting rate (α_F_, y-axis). Plots are organized based on the intrinsic rate of increase (*r*), per capita predation pressure (*H*), and the initial abundance of the rare prey. Columns in (A-D) are results when the rare prey was 10%, 50%, and 90% as abundant as the common prey. Black cells in (A) and (C) denote the extinction of both signals, red cells denote the extinction of only the rare signal, and blue cells denote the persistence of both signals. In (B) and (D), green colors indicate negative selection on the rare prey, brown colors indicate positive selection, and gray color denotes a selection value of zero. (A) and (B) are when prey have separate carrying capacities, and (B) and (D) are when they share the same carrying capacity. When prey have separate carrying capacities, low asymptotic attack probability is conducive to the persistence of the rare signal, whereas intermediate asymptotic attack probabilities have the same effect when prey share a carrying capacity. In all scenarios, a sufficiently low forgetting rate is the most limiting condition for local warning signal diversity.

**Figure 2.**
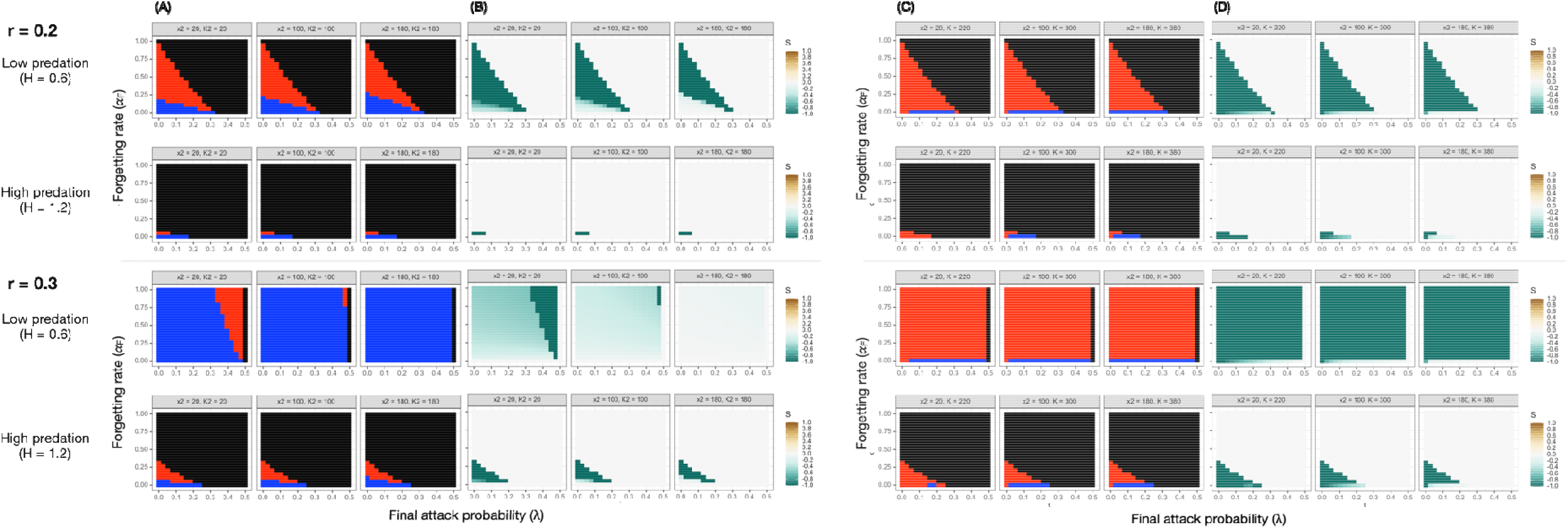
Prey persistence (A, C) and corresponding selection on the rare prey (B, D), with respect to intrinsic rate of increase (*r*), per capita predation pressure (*H*), and the initial abundance of the rare prey when there is generalization (G = 0.5, see main text). Generalization facilitates local warning signal diversity in all scenarios, especially when the ecological conditions are already favorable (e.g., high intrinsic rate of increase and low per capita predation pressure). The organization of the plots and coloring of the cells are the same as in Figure 1.

Notably, across all simulations the persistence of the rare signal did not always correspond to positive selection values, especially when prey shared a carrying capacity (Figs. 1, 2, S2, S3, S5 and S6). This demonstrates that although rare prey can be less successful in terms of relative change in abundance, positive FDS alone does not predict whether warning signal diversity can occur in communities. The individual-based model produced qualitatively similar results when prey did not compete (Fig. 3A). In the presence of prey competition, however, it required even stronger predation to offset the competitive advantage of the common prey compared to the population-level model, because reproduction only happened at fixed intervals instead of occurring at all times. This was reflected in the fact that warning signal diversity occurred under very high final attack probabilities and non-zero forgetting rates in Fig 3B.

**Figure 3.**
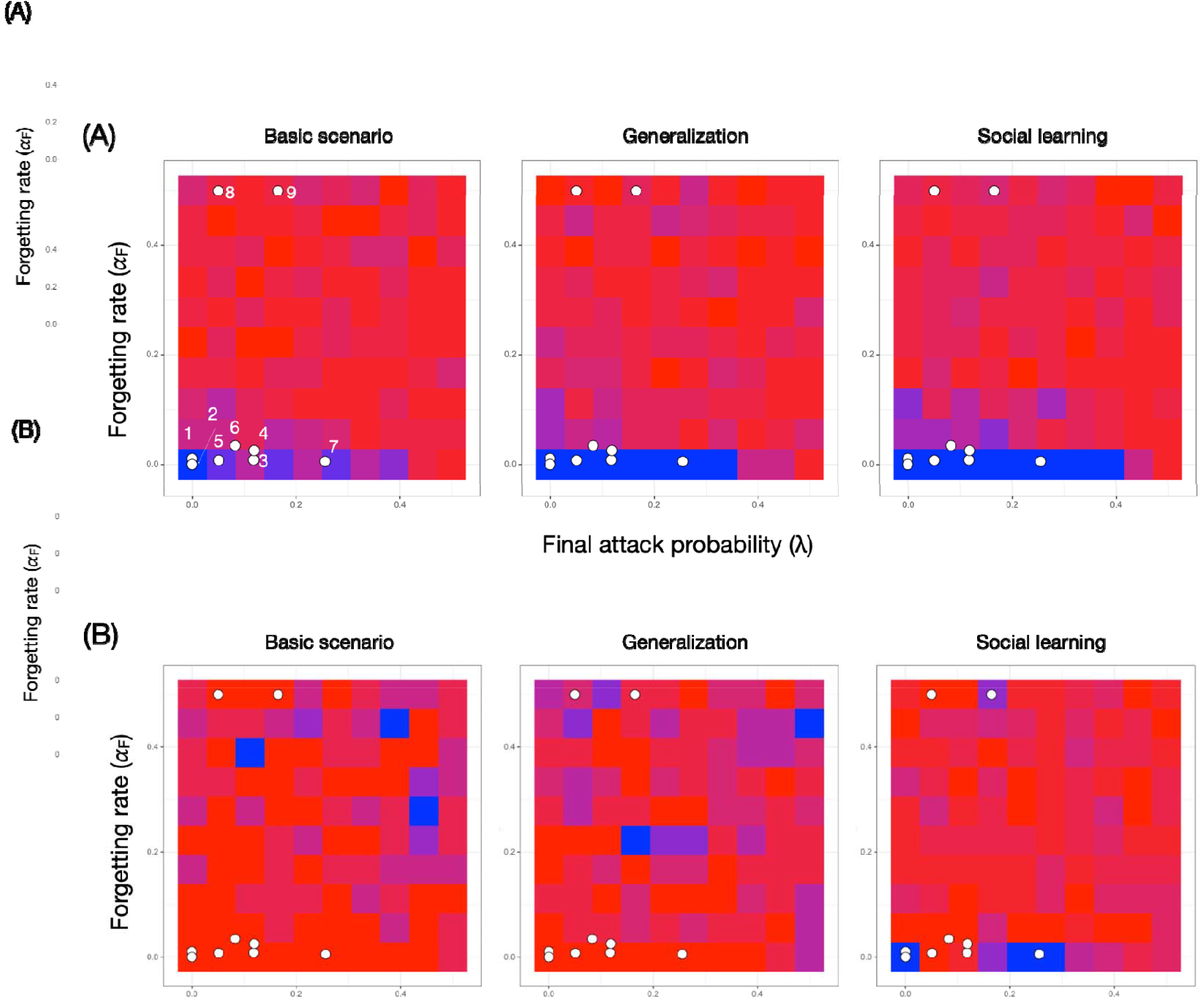
Prey persistence with respect to final attack probability and forgetting rate from the individual-based simulations when prey have separate carrying capacities (A) or when they share the same carrying capacity (B). Within (A) and (B), plots are grouped to show results from the basic scenario, with generalization, and with social learning under *r* = 0.2 (top panel) and 0.3 (bottom panel). The columns within each scenario are when the rare prey was 10%, 50%, and 90% as abundant as the common prey. The coloring of the cells are the same as in Figures 1 and 2.

Both generalization and social learning increased the likelihood of warning signal diversity (Figs. 2 and 3). To see if the parameter value combinations favorable for warning signal diversity corresponded well to what has been observed in real predators, nine pairs of final attack probabilities and forgetting rates from six species (one spider, one fish, one lizard, two birds, and one mammal) were mapped onto the parameter space (Table S1 and Fig. 4). The focus here was not the species identity *per se*, but the combination of their final attack probabilities and forgetting rates. There were only nine data points because forgetting rates were rarely quantified in avoidance learning studies. Even with limited number of cases, the combination of avoidance level and forgetting rate from most of the predators were at the appropriate locations in the parameter space, suggesting that predators from a wide range of taxa could potentially maintain local warning signal diversity. Empirical values of generalization and social learning efficacy are listed in Table S2.

**Figure 4.** Examining the potential of predators in maintaining local warning signal diversity by mapping empirically measured final attack probability and forgetting rate onto the parameter space (white points): 1. *Planigale maculata* (91), 2. *Habronattus pyrrithrix* (54), 3-5. *Parus major* (92), 6. *Chalcides sexlineatus* (93), 7. *Gallus gallus domesticus* (94), 8-9. *Thalassoma bifasciatum* (56). The plots here are from simulations with r = 0.3 and the relative abundance of the rare prey being 10%. (A) Prey have separate carrying capacities. (B) Prey share the same carrying capacity.

### Determinants of final attack probabilities

Compared with bitter prey, there was very strong evidence that toxic prey led to a higher probability of complete avoidance (i.e., an attack probability of zero), but being toxic did not reduce non-zero attack probabilities (Table 2 and Fig. 5A). There was also no evidence that increasing bitterness led to lower attack probabilities (z = −0.23, P = 0.82, Fig. 5B). There was very strong evidence that predators avoided unprofitable prey more when profitable prey were also present (Table 2 and Fig. 5C). Finally, there was no evidence that predators avoided unprofitable prey differently when the feature distinguishing prey types involved only color, only pattern, or color plus size or odor (Table 2 and Fig. 5D). However, when the signals or cues differed in both color and pattern, there was very strong evidence that predators avoided unprofitable prey less, suggesting that signals containing complex visual information might be less effective for predator education, at least for the four avian species examined (Table 2 and Fig. 5D).

**Figure 5.**
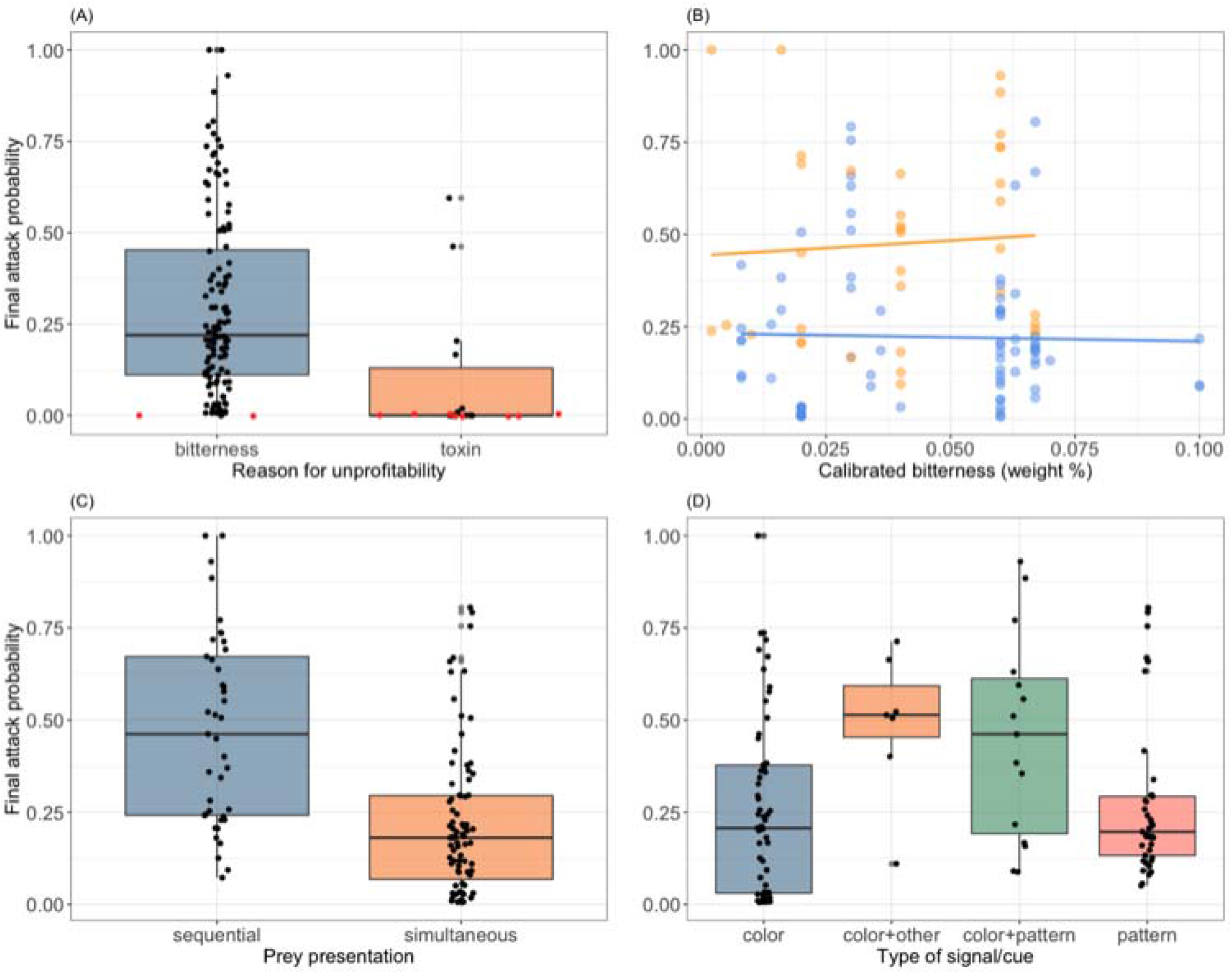
Factors that influenced predators’ final attack probabilities towards unprofitable prey. (A) Reason for unprofitability, (B) level of bitterness, (C) manner of prey encounter, and (D) type of signal or cue. Red dots in (A) are attack probabilities that equal zero. Yellow dots and line in (B) are when predators encounter profitable and unprofitable prey sequentially, and blue dots and line are when predators encounter the two prey types simultaneously. Data were from (31, 32, 46–49, 53, 86, 89, 92, 94–121)

## Discussion

### Local warning signal diversity is not paradoxical, and the detection of positive FDS alone is not a reliable indicator of its possibility without knowing predator forgetting rate

When predators can retain learned avoidance for long enough, the persistence of rare signals in communities can be achieved through balanced predation and reproduction and is entirely nonparadoxical. Notably, high prey unprofitability only facilitates warning signal diversity when prey displaying different warning signals do not compete with one another. This condition was also thought to facilitate the evolution of aposematism from crypsis in unprofitable prey (36). When prey do compete, warning signal diversity can only result when prey elicit moderate avoidance. Empirical learning data also suggest that natural predators from a range of major taxa may have sufficiently low forgetting rates and appropriate levels of prey avoidance to allow for local warning signal diversity. However, local warning signal diversity is unlikely to occur if the rare signal is less than 10% in relative abundance with prey competition and without generalization or social learning. Clearly, this value depends on the prey intrinsic rate of increase and should not be taken as a hard threshold. Even though the relative abundance of stably coexisting warning signals has only been quantified for a small number of systems, it appears to be generally true that the rare signals often constitute at least ∼ 10% of the population (37–39). The result may also help further explain the observation that warning signal diversity is often the result of secondary contact following dispersal (40–42), in which neither signal is likely to be astronomically dominant in number.

The fact that the rare signal can persist despite negative values of selection might appear counterintuitive at first glance. The expectation that a phenotype granting higher relative fitness will ultimately eliminate other phenotypes is based on the premise that relative fitness translates to relative abundance in every generation, or at least regularly enough. This may not be true for prey whose survival is determined by learned avoidance from sufficiently unforgetful predators, especially when they have separate carrying capacities. Even though selection is calculated using the initial and final prey abundance, in effect it only captures the difference in relative fitness between onset of predator learning and the steady-state. Negative values of selection on rare prey therefore only reflects the fact that their relative abundance decreased more compared to the common prey up until the steady-state, but not beyond. The important implication of this result is that empirically detected positive FDS alone, especially those examined when predators are still learning (e.g., ref. 37), does not reliably inform the fate of rarer warning signals within a community.

The diversity and convergence of warning signals at the regional and local scales has been thought to reflect a shifting balance involving local drift (phase I), local selection (phase II), and the spread of competing, locally adaptive peaks from local to regional scales (phase III)(5, 25). Results in this study offer further ecological insights into the shifting balance process. First, generalization and especially social learning can further flatten the “adaptive trough” for rare warning signals (Fig. 1 in ref. 5). This can facilitate the establishment of new waring phenotypes locally through processes such as kin founding, especially if individuals with the new warning signals have a fitness advantage, for example, from being more unprofitable or more conspicuous. When all prey are equally unprofitable, the facilitative effects of generalization and social learning are only apparent when the prey intrinsic rate of increase is sufficiently high. The fact that positive FDS does not necessarily lead to the elimination of rarer warning signals also points to an alternative outcome during phase III of the shifting balance, in which two adaptive peaks meet without one replacing the other.

Aside from predator forgetting, positive FDS can result when predator turnover rate is comparable to prey generation time, such that prey face naïve predators in every generation. Even though the simulations did not directly model predator turnover, the erosion of learned avoidance over time can be simulated as a consequence of predator turnover in a community that is similar to forgetting of individual predators. For example, if each predator in the model is treated as a group that contains multiple individuals, the same process in which attack probabilities increase towards 0.5 over time can represent the replacement of experienced predator individuals with naïve ones in each predator group. The only modification required for simulating predator turnover in the model is that the erosion of learned avoidance will occur at each time step regardless of whether predators attack the focal prey. In contrast with predator forgetting, predator turnover is guaranteed to take place at each time step and can therefore have a stronger limiting effect on the likelihood of local warning signal diversity. Since most Müllerian prey are insects and frogs, a substantial turnover of predators within one prey generation may be unlikely (e.g., one month for butterflies and one year for poison frogs, refs. 43, 44). For Müllerian prey with longer generation time, such as the putative mimicry rings between some mustelid, viverrid, and herpestid mammals (45), significant predator turnover could occur between prey generations and thus might play a more pivotal role.

### Variation in final attack probabilities and the implications for local warning signal diversity

Even though data in the meta-analyses were conducted on a small number of birds under laboratory settings, findings from the meta-analysis are broadly informative as the experiments were all designed to capture the essence of predator learning. The analysis revealed that being toxic can be more effective than unpalatability for dissuading some predators, resulting in complete avoidance. However, for predators that are not completely deterred by toxicity, either because of a higher toxicity tolerance or more urgent energetic needs, being toxic did not appear to offer more protection than mere bitterness. Studies that presented toxic food to predators either used naturally toxic prey (46, 47) or prepared artificial food items with extracted prey toxin (48). These prey have chemical defenses that dissuade rather than severely harm predators, which is how defense is expected to function in Müllerian systems. It is therefore reasonable to extrapolate this result to how predators might avoid Müllerian prey in general. In addition, increasing bitterness does not appear to be more effective in eliciting higher avoidance, at least within the range of bitterness in the analysis (weight percentage 0.002 – 0.1). It is possible that the 50-fold difference in the effective concentration of bitter chemicals does not result in corresponding changes in bitter perception in birds. Alternatively, this result could indicate that birds respond to prey bitterness in a uniform fashion, despite continuous bitterness perception. Data on taste sensory physiology, coupled with controlled tests of taste response, would help differentiate these two hypotheses. All in all, the meta-analysis suggests that Müllerian systems with toxic prey could have higher potential for local warning signal diversity, but only if the toxins cause complete avoidance. On the other hand, local warning signal diversity is equally likely to occur in Müllerian systems with highly distasteful species than in those whose prey are less obnoxious in taste. Evasiveness is another important cause of prey unprofitability but is not included in the meta-analysis because only one study has quantified attack probabilities towards these prey; evasive prey elicited a lower level of avoidance but a stronger generalization effect than chemically defense (49). Examining predator learning in more such systems (e.g., the Amazonian *Morpho* butterflies, refs. 50, 51) will be important for further testing whether local warning signal diversity is more or less likely to occur in these evasive Müllerian systems.

Predators avoid unprofitable prey more when profitable alternatives are present in the environment. Even though this result comes from laboratory experiments, it has important implications for predator-prey interactions in natural communities. The simultaneous and sequential prey presentations in laboratory experiments represent two extremes of the prey-encounter continuum. In the field, the foraging habits of predators and the density and distribution of prey can both determine the manner of predator-prey interactions. Sit-and-wait predators are more likely to encounter prey in a sequential manner than widely ranging predators and therefore might avoid unprofitable prey to a lesser extent. Similarly, predators are more likely to encounter prey sequentially in habitats where prey density is low and/or where prey do not aggregate. Under these conditions, predators are also expected to exhibit lower avoidance towards unprofitable prey. The meta-analysis suggests that, all else being equal, local warning signal diversity is more likely to occur in communities when main predators actively search for prey and when prey are abundant and/or have a concentrated distribution, provided that prey do not compete. These predictions have never been tested. In addition to predator forgetting rate, a comparative analysis examining the relationship between the occurrence of local warning signal diversity, predators’ foraging habits, and prey distribution would offer valuable insight.

The nature of the warning signal can also affect predators’ final attack probabilities; predators avoided unprofitable prey more when they differed from profitable prey in only color or pattern. Furthermore, colors also differ in their educational efficacy as learning cues (52–57). Based on this finding, local warning diversity might be more easily achieved in communities without considerable competition between prey species if profitable prey have simple visual appearance and could be differentiated from unprofitable prey based on color or pattern alone. This does not mean that unprofitable prey cannot possess visually complex warning signals, as long as predators can prioritize on color or pattern for foraging decision-making (unequal feature salience or overshadow effects, refs. 58–63). However, it is worth noting that this finding was based on results from a small number of species and might have limited generality. Particularly, we know next to nothing about avoidance learning abilities in tropical birds, which are the main predators of the most classic Müllerian prey (e.g., refs. 64–66). Quantifying learning and forgetting in tropical avian predators can be challenging but would provide crucial information on the maintenance of local warning signal diversity in some of the most remarkable Müllerian systems.

### Testing the Unforgetful Predator Hypothesis in natural communities

All key parameters of the unforgetful predator hypothesis - final attack probability, forgetting rate, strength of generalization, and the efficacy of social learning - can be measured with learning experiments. Usually, these parameters are measured in a laboratory setting, which allows data collection from individual predators and is particularly useful in assessing the variation in learning parameters within and among individuals. In communities where individual predators can be identified and their learning parameters estimated, the model can be easily modified to properly reflect the heterogeneity in predator behavior and cognition. From available data, it appears that predators with simpler nervous systems (e.g., spiders) can have forgetting rates comparable to those of birds and mammals. This suggests that predators with high potential for maintaining local warning signal diversity are not limited to more derived taxa and may be more common than expected.

When collecting learning data from individual predators are not feasible, field learning experiments represent a suitable alternative. Field learning experiments present artificial prey to all potential predators in the community instead of targeting specific predator individuals or species (e.g., refs. 67–74). Final attack probabilities, forgetting rates, and the strength of generalization can be derived from attack rates in the same way as the laboratory experiments. The key difference is that learning parameters obtained in these experiments can be considered as weighted averages over all predator species. These field learning experiments can be very useful in testing the Unforgetful Predator Hypothesis in natural communities with highly heterogeneous predator compositions. The efficacy of social learning can also be estimated in field learning experiments by comparing attack rates from predators that learn from watch video playbacks and those that learn directly. So far, neither the forgetting rate nor the efficacy of social learning has been quantified with field experiments. Filling this knowledge gap is crucial for more broadly understanding the role of predator learning in the maintenance of waring signal diversity in natural communities.

In addition to key predator learning parameters, population genetic/genomic data can help further elucidate whether the presence of rare warning signals in a community is in fact ecological persistence maintained by unforgetful predators. If the genetic diversity among individuals displaying the rare signal in the focal community is lower than the adjacent “source” populations, this would indicate that prey with rare signals represent a founder population that receives limited immigration from the source populations. Conversely, if the genetic diversity is comparable between the focal and source populations this would suggest the focal population might have been experiencing rapid turnovers between predation-induced deaths and immigration that may not be stable over longer time scales. Demographic analyses indicating a significant reduction in effective population size for prey bearing rare signals can provide another line of evidence supporting warning signal diversity is being maintained in the focal community (75). Lastly, our findings that warning signal diversity requires different levels of predator avoidance and forgetting rates in different ecological environments complement results from (76), which showed that variation in predator behavior over regional scales could underlie warning signal polymorphism within species.

## Materials and Methods

All simulations and statistical analyses were performed in R (version 4.2.2, R Core Team).

### Simulating predator-prey interactions and predator learning

I simulated population dynamics of prey displaying different warning signals by constructing both population-level and individual-based models. The population-level model offered deterministic insights into the ecological dynamics of prey populations as influenced by predator learning, forgetting, and reproduction. Since predator learning is fundamentally an individual-level process, I accompanied the population-level model with the individual-based model, which better encapsulated the individual aspects of predator-prey interactions and was better suited for modeling the dependence of learning experience between predators. In both types of models, I considered two ecological scenarios regarding whether prey displaying distinct warning signals did not compete and therefore had separate carrying capacities, did compete and shared the same carrying capacity. When prey had separate carrying capacities, I further considered two situations; one in which the abundance of the rare prey was constrained to never exceed its initial value, and the other in which the rare prey can potentially reach the same abundance as the common prey (Table 1). I considered similar situations when prey shared the same carrying capacity; one in which the shared carrying capacity equaled to the sum of initial prey abundances, and the other in which the carrying capacity equaled to twice the abundance of the common prey (Table 1).

**Table 1.** Definitions and values of state variables and parameters in the population-level model.

| Variable/parameter | Definition | Value in the models |
| --- | --- | --- |
| Common to all scenarios |  |  |
| $X_{0,1}$ | Initial abundance of the common prey | 200 |
| $X_{0,2}$ | Initial abundance of the rare prey | 20, 100, 180 |
| $r$ | Intrinsic rate of increase | 0.2, 0.3 |
| $H$ | Per capita predation pressure | 0.6, 1.2 |
| $\lambda$ | Final attack probability | 0 – 0.5 |
| $\alpha_F$ | Forgetting rate | 0 – 1 |
| $G$ | Carrying over learning effect due to generalization | 0, 0.5 |
| Separate prey carrying capacities |  |  |
| $K_1$ | Carrying capacity of the common prey | 200 |
| $K_2$ | Carrying capacity of the rare prey | 20, 100, 180<sup>a</sup><br>200<sup>b</sup> |
| Shared carrying capacity for both prey |  |  |
| $K$ | Shared carrying capacity | 220, 300, 380<sup>c</sup><br>400<sup>d</sup> |

<sup>a</sup>When the abundance of the rare prey was constrained to the initial abundance.
<sup>b</sup>When the rare prey could potentially be equally abundant as the common prey.
<sup>c</sup>When the shared carrying capacity equaled to the sum of initial prey abundances.
<sup>d</sup>When the shared carrying capacity equaled to twice the initial abundance of the common prey.

**Table 2.**
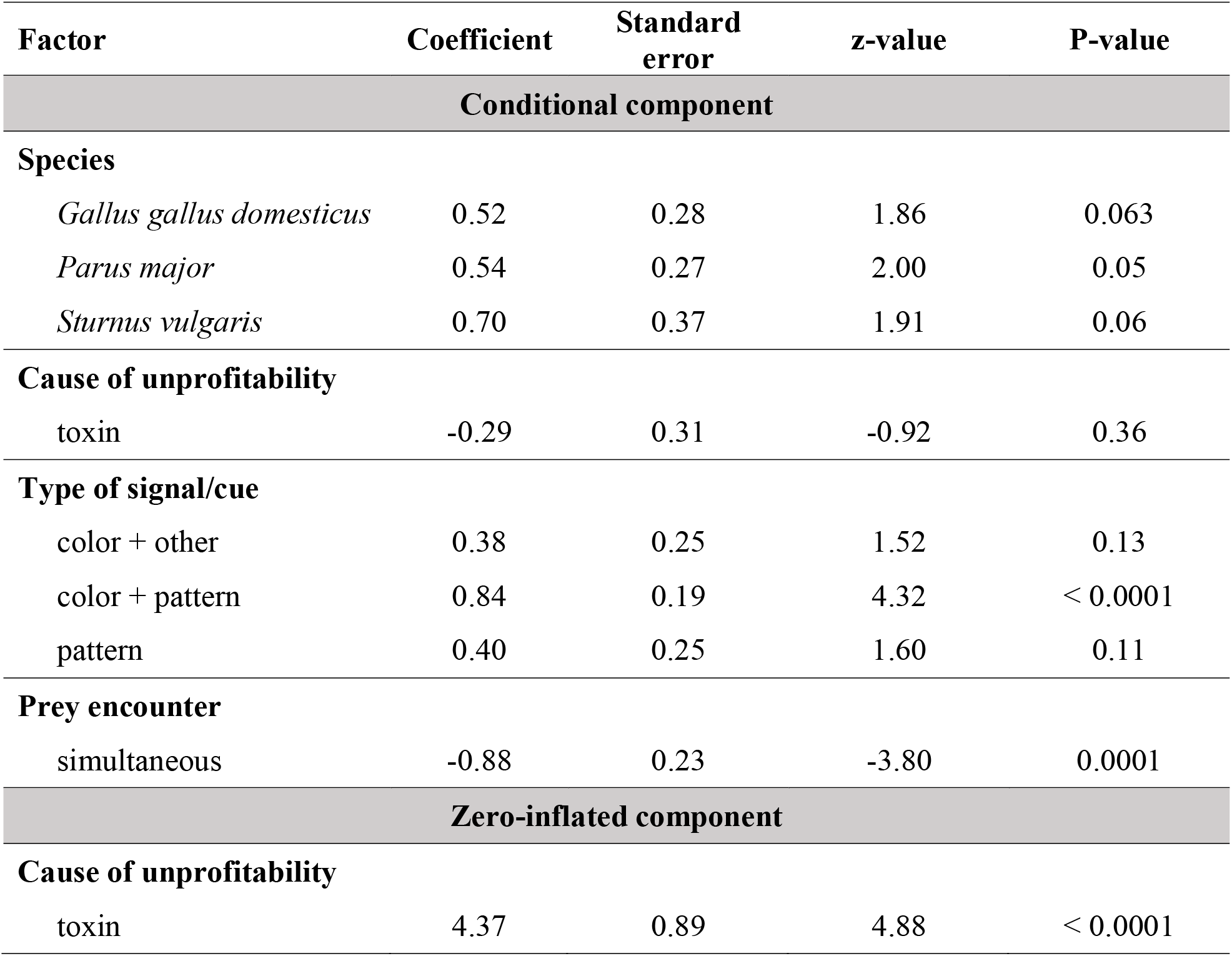
Statistical results of the zero-inflated gamma regression examining the cause of variation in predators’ final attack probabilities towards unprofitable prey.

I did not consider spatial segregation or habitat structuring in all models as the intention here was to focus on explaining local warning diversity with predator learning alone. As the coexistence of distinct warning signals in well studied Müllerian mimetic systems is mostly the result of secondary contact (40–42), the model also did not consider the appearance of novel warning signals by mutations within populations.

### The population-level model

In a fully-mixed community of predators and prey with learning occurring simultaneously for all predators, the instantaneous change in prey population size can be described by the following equations when prey had separate carrying capacities:

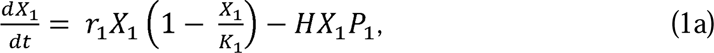

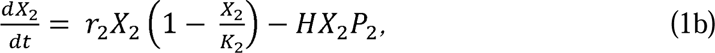

where *X* is prey population size, *r* is the intrinsic rate of increase for each prey type and *K* is prey carrying capacity. In all simulations, prey one are more common than prey two. When all prey shared a carrying capacity (K), instantaneous changes in population size become

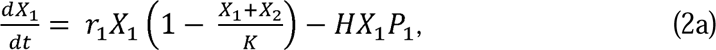

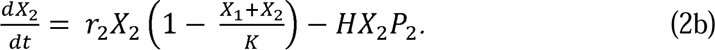

The first term in these pairs of equations describes instantaneous population increase due to reproduction, and the second term describes instantaneous decrease in population size due to predation. The term *H* integrates information on predator abundance and foraging efficiency (e.g., (77) and could be considered as the per capita predation pressure unrelated to predator learning. *P* is the attack probability towards prey. However, the attack probability *P* also changes with time as a dynamic tugging between learning and forgetting. Based on the Pavlovian learning framework (14–20), change in attack probability due to learning (Δ*P_L_*) can be defined as

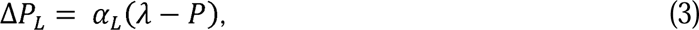

where λ is the asymptotic attack probability ranging from 0 to 0.5 for unprofitable prey, and α_L_ is the learning rate, defined as a function of λ

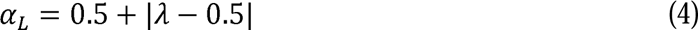

At the same time, attack probability towards unprofitable prey can also change due to forgetting (Δ*P_F_*).

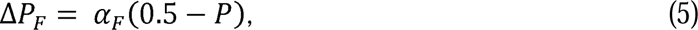

where α_F_ is the forgetting rate, with values ranging from 0 (no forgetting) to 1 (attack probability returning from the current value to 0.5 in one single time step).

Taken together, the instantaneous changes in predators’ attack probabilities towards the two prey types can be described with the following pair of equations.

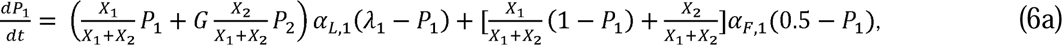

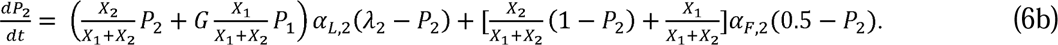

These equations describe changes in attack probabilities as a joint consequence of learning when predators encounter and attack the focal prey and forgetting when they do not. The term *G* accounts for the carrying over learning effect from cognitive generalization. When *G* equals zero (i.e., no generalization), predators can only modify their attack probabilities towards the focal prey from first-hand learning.

I examined the equilibrium population sizes for the two prey with different starting population sizes and values for *r, K, H,* λ, α_F_, and *G* (Table 1) by simultaneously solving equations 1a, 1b, (or 2a and 2b), 6a, and 6b using the *ode* function in the R package deSolve (78). Population sizes were solved for 10,000 steps with a 0.1 increment (a total of 100,000 values). To be ecologically realistic about prey’s intrinsic rate of increase, I used *r* values of 0.2 and 0.3, which encompassed the range of *r* commonly observed in insects and some mammals (79–85). I used *H* values of 0.6 and 1.2 to represent low- and high-predation environments. These *H* values were selected based on preliminary simulations to produce sufficiently different yet ecologically sensible outcomes (e.g., extinction of only the rare prey can be observed). I verified that the equilibrium has been reached for all simulations by confirming that the coefficient of variation for the last 1,000 abundance values was zero for both prey. I also calculated selection (*S*) on prey displaying the rare signal based on proportional changes in abundance relative to the common prey as

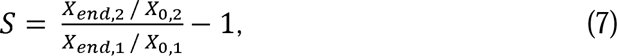

where X_0_ and X_end_ denote the initial and final prey abundance. If both prey go extinct, *S* was assigned the value of zero.

### The individual-based model

I constructed individual-based models with discrete time steps and simulated interactions between predators and aposematic prey in one single habitat, while considering the effects of generalization and social learning. There were 20 predator individuals and two prey types in the simulated community. Avoidance learning here also followed the Pavlovian framework (eq 3 and 4, (14–20). I modelled forgetting here as simply a fixed increase (i.e., forgetting rate) of attack probability whenever the predators did not attack the focal prey, with the attack probability not exceeding 0.5 after the increase. The values of forgetting rate in individual-based models therefore ranged from 0 to 0.5. This was different from the Pavlovian framework but had the important advantage of allowing the use of empirically measured forgetting rate for model validation (see *Testing the unforgetful predator hypothesis with empirical learning data* below).

Each simulation was run for 50,000 steps, and in each time step a predator was randomly selected to be the forager. The forager would encounter prey based on relative abundance and then decide whether to attack the prey based on the current attack probability towards that prey type. Throughout the simulation, predators continued to modify their attack probabilities towards the two prey types following the Pavlovian learning rules. The attack probability towards the familiar prey was set at its final asymptotic value at the beginning of the simulation, reflecting the fact that predators have completed learning. The attack probability towards the new prey was set at 0.5 at the onset of the simulation, signifying predator naivete. Reproduction occurred every 100 steps in all simulations. No predation occurred during reproduction and both prey types replenished their populations following a logistic growth function as described in equations 1 and 2, depending on whether the two prey had separate carrying capacities. For each simulation, the parameter space was a 10 by 10 grid defined by the final attack probability and forgetting rate, both of which ranged from 0 to 0.5. I ran 10 replicate simulations for each combination of parameter values. Abundance of both prey types were recorded at the end of each simulation. Selection on the rare waring signal was calculated with eq 7.

The above procedure described the basic scenario, in which avoidance learning for the rare and the common prey occurred independently – predator learning with one prey type did not alter their attack probabilities towards the other prey type. Each predator also learned independently. To model the effect of generalization, I introduced a parameter *G* as in population-level models, such that learning occurring at time *t* was transferable between prey types:

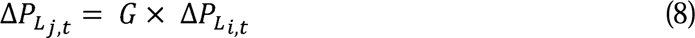

In addition,

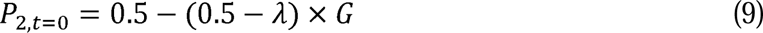

The equation above reflected the fact that the initial attack probability towards the rare prey will be reduced based on its similarity to the common prey. When *G* equaled one, the attack probability towards the rare prey would already be identical to the final attack probability at the onset of the simulation, and learning was completely transferable between prey types. A *G* equaled zero was equivalent to the basic scenario described above. In simulations with generalization, *G* was set to be 0.5.

In separate sets of simulations, I incorporated social learning similar to (33) by introducing two parameters. The first parameter *b* determined the percentage of non-forager individuals that were able to observe the prey interaction of the forager. Larger *b* values meant that more predators could benefit from social learning. The other parameter *q* described the efficacy of social learning. Similar to how the parameter *G* functioned, *q* was a percentage denoting a “discount” in learning effect relative to first-hand experience, such that

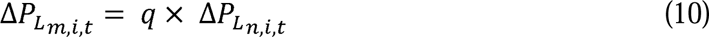

At time step *t*, predators *m* observed the foraging outcome between the forager *n* and prey type *i.* If *q* equaled one, social learning would cause the same decrease in attack probability for predator *m* as it would from direct learning. If *q* equaled zero, there was no social learning. In simulations with social learning, both *b* and *q* were set to be 0.5.

### Testing the Unforgetful Predator Hypothesis with empirical learning data

#### Extracting final attack probabilities and forgetting rates from published studies

I performed literature search using keyword combinations (topic = *aposemat\** or defen* or *avoid\** + topic = learning) in the Web of Science database, yielding a total of 387 articles. I then screened for studies that quantified attack probability in the context of avoidance foraging with focal animals under normal conditions (e.g., not injured, not sick, not under high stress, not surgically or genetically modified). This resulted in 112 articles from which I extracted attack probabilities. These studies all followed a standard experimental design, in which animals were presented with prey in a series of learning trials. I calculated predators’ attack probabilities towards the unprofitable prey from the last trial as the final attack probability.

Studies included in the meta-analysis all presented predators with equal number of unprofitable experimental prey and profitable control prey that differed consistently in certain aspect of appearance. The locations of prey are randomized in the test arena such that the predators are expected to encounter equal number of each prey type. The experiments ended when the predator attacked a predetermined total number of prey, regardless of type. Even though the ratio between prey attacked and prey of the same type presented is widely reported as predation risk or mortality rate, this ratio does not represent the attack probability towards that prey type. Instead, the attack probability was calculated as the ratio between the number of focal prey attacked and the number of control prey attacked. The rationale behind this calculation can be explained using an analogy of tossing two separate biased coins for equal amount of times *N* until a predetermined number of heads (i.e., attack) is reached. Under this condition, the number of heads from coin A (n_A_) and B (n_B_) will directly reflect the degree of bias of each coin, such that

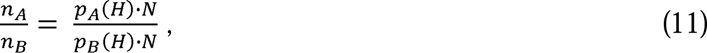

where *p_A_* (*H*) and *p_B_* (*H*) denote the probability of heads from coin A and B, respectively. It is clear that this degree of bias cannot be estimated for coin A unless the degree of bias is known for at coin B, and *vice versa*. In the learning experiments examined, it is reasonable to assume that the attack probability towards profitable control prey (i.e., *p_B_* (*H*) is very close to 1, thereby allowing us to infer predators’ attack probability towards the unprofitable prey (i.e., *p_A_* (*H*)(Fig. S1).

In studies that measured predator forgetting, forgetting rate was calculated as the difference in attack probabilities between trials immediately before and after the forgetting period, divided by the duration of that period. These forgetting rates were then scaled, when necessary, to be percent changes per day (see *Mapping empirical learning data onto the model parameter space*).

#### Quantifying the effect of generalization and social learning

To examine generalization, studies allowed animals to complete learning towards a prey type and then presented the animals with a new prey type that bore similarities to the familiar prey.

I calculated the magnitude of generalization as

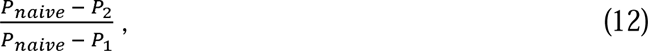

where *P_naive_* was the attack probability towards the first prey type at the beginning of the experiment, *P_1_* the attack probability towards the first prey type at the end of the experiment, and *P_2_*the attack probability towards the second prey. The denominator described changes in attack probability towards the first prey due to direct learning, and the numerator quantified the effect of generalization on attack probability towards a second prey type. This ratio scaled the strength of generalization by the learning effect on the first prey type, making it analogous to how the parameter *G* functioned in the model.

The efficacy of social learning was examined in three studies from the literature search, yet there appeared to be two different types of learning outcomes. In two studies, social learning caused the predators to avoid the same unprofitable prey even more than they would from first-hand learning; the final attack probabilities towards the same unprofitable prey were lower in socially informed predators than predators that learned from direct experiences (31, 32). For these studies, the efficacy of social learning was the difference in final attack probabilities between direct and social learners, divided by the final attack probability of direct learners. In one study, social learning did reduce attack probabilities towards unprofitable prey, though not to the same extent as direct learning (86). I calculated the efficacy of social learning from this study in the same way as I did with generalization:

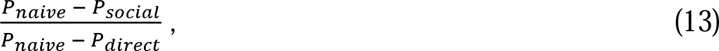

where *P_naive_* was defined as in eq. 12, *P_social_* was the attack probability towards the focal prey from social learners, and *P_direct_* was that from first-hand learners. Similarly, these calculations scaled the efficacy of social learning by the effect of direct learning, making it analogous to how the parameter *q* functioned in the simulations.

#### Mapping empirical learning data onto the model parameter space

For predators whose final attack probabilities and forgetting rates towards unprofitable prey are quantified, their data can be mapped onto the parameter space of the model to evaluate their potential for permitting local warning signal diversity. Empirically measured final attack probability was immediately appropriate for the parameter space in the simulations. Mapping empirical forgetting rates onto the parameter space required one extra step. In the model, each time step equaled 0.01 prey generation time. Using a prey generation time of 30 days, each time step in the model was then equivalent to 0.3 days. An empirical forgetting rate of 0.1 per day would thus equal to a value of 0.03 in the model. In general, any empirically measured forgetting rate could be scaled to unit per day and converted to values appropriate for the parameter space by multiplying the per day value with 0.01 focal prey generation time.

### Meta-analysis of the variation in final attack probability

To obtain a sufficiently balanced dataset, I focused on 39 of the 112 previously selected studies for the meta-analysis, yielding a total of 134 final attack probability measurements (Supplementary Data). These studies shared enough similarities in experimental design while varying in crucial aspects, allowing for meaningful analyses. The focal predators were either the great tit (*Parus major*), the blue tit (*Cyanistes caeruleus*), the European starling (*Sturnus vulgaris*), or the domestic chicken (*Gallus gallus domesticus*). In addition, animals were presented with both control and unprofitable artificial prey within each learning trial, either simultaneously or sequentially. The reason for prey unprofitability was either bitterness (using harmless, bitter chemicals) or chemical defense (using prey extracts), and the feature that consistently differentiated the two prey types was either color alone, pattern alone (i.e., the distribution of colors or the use of different geometric symbols), both color and pattern, or color and size or odor.

To perform statistical analyses, I used individual measurements of final attack probability as the response variable, with species and the aforementioned three factors as predictors. Individual study was originally included as a random-effect factor to account for potential nonindependence of data points within studies. However, doing so resulted in model overfitting, and I excluded study from the statistical model. I also did not consider the phylogenetic relationship between species since the number of species was small. Although final attack probabilities were originally proportions derived from count data, the nature of these proportions was different from that derived from binomial trials. I therefore treated the final attack probabilities as continuous data modeled with a gamma distribution when constructing subsequent statistical models. The choice of gamma over beta distribution was because the latter did not allow values equaled to one, which existed in the dataset. To examine the causes of variation in final attack probability, I first performed a zero-inflated gamma regression using the *glmmTMB* package (87). In the model, the reason for unprofitability was specified as the factor causing zero-inflation, and dispersion parameters were allowed to vary among levels within each fixed-effect factor. I did not consider interactions among predictors, as the dataset were not sufficiently replicated across all factor combinations. I performed the statistical test using the package *glmmTMB*. All models were validated with residual diagnostics using the *DHARMa* package (88) before interpretation.

To additionally test whether the effective concentration of the bitter chemical was associated with variation in final attack probabilities, I focused on 35 of the 39 studies that used bitter experimental prey. The majority of the studies used quinine derivatives, whereas four studies used denatonium benzoate (Supplementary Data). A study reported that 2% quinine sulfate and 0.1% denatonium benzoate (weight percentages) elicited comparable aversion in domestic chicks (89), signifying a 20-fold difference in effective concentration between the two. I therefore converted the concentrations of denatonium benzoate to equivalent concentrations of quinine derivatives by multiplying them by 20. To test whether an increase in the concentration of bitter compounds caused lower attack probabilities, I performed a gamma regression with manner of prey encounter, the standardized concentration of bitter compounds, and the interaction between the two as fixed-effect predictors. When reporting results, I used the language of evidence instead of significance dichotomy (90).

## Statement of authorship

CYK conceived the study, performed computer simulations, literature search and meta-analysis, and wrote the manuscript

## Data accessibility statement

The dataset and R codes for the simulation and meta-analysis can be accessed at https://github.com/thekuolab/warning-signal-diversity

## Acknowledgements and funding sources

I thank Po-Ju Ke and Thomas Sherratt for their invaluable insights that helped improve this paper. The author also thanks Yi Sun and the McMillan lab at the Smithsonian Tropical Research Institute, and two anonymous reviewers for their helpful comments on earlier drafts of this manuscript. This study was supported by the Kaohsiung Medical University Research Foundation (grant number: KMU-Q111001 and KMU-Q112001).

## Notes

https://github.com/thekuolab/warning-signal-diversity

### Competing Interest Statement

The authors have declared no competing interest.

### Summary of Updates

The Introduction and the Discussion sections have been significantly revised. I also included new continuous-time simulations to accompany the individual-based models, which expanded the scope of the analyses. Some methodological details that were in the Supplementary Materials are now in the main text.

