## Supplementary Tables and Figures for "Resolving the paradox of local warning signal diversity with predator learning"

**Supplementary Figures and Tables**

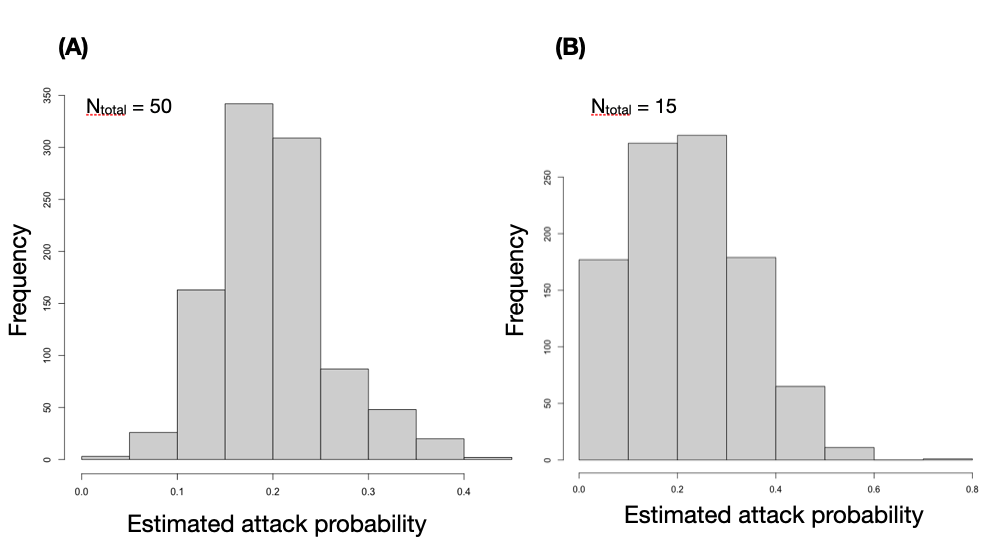

**Figure S1.** Estimations of attack probability using equation (1) from 1000 replicate simulations, in which predators were presented with profitable (attack probability = 1) and unprofitable prey (attack probability = 0.2). A simulation ended when the predator had attacked 50 (A) or 15 prey (B). The simulation results showed that the estimated attack probabilities had a central tendency around the true value of 0.2, but the variation was smaller when predators were allowed to attack a larger number of prey before simulation ended.

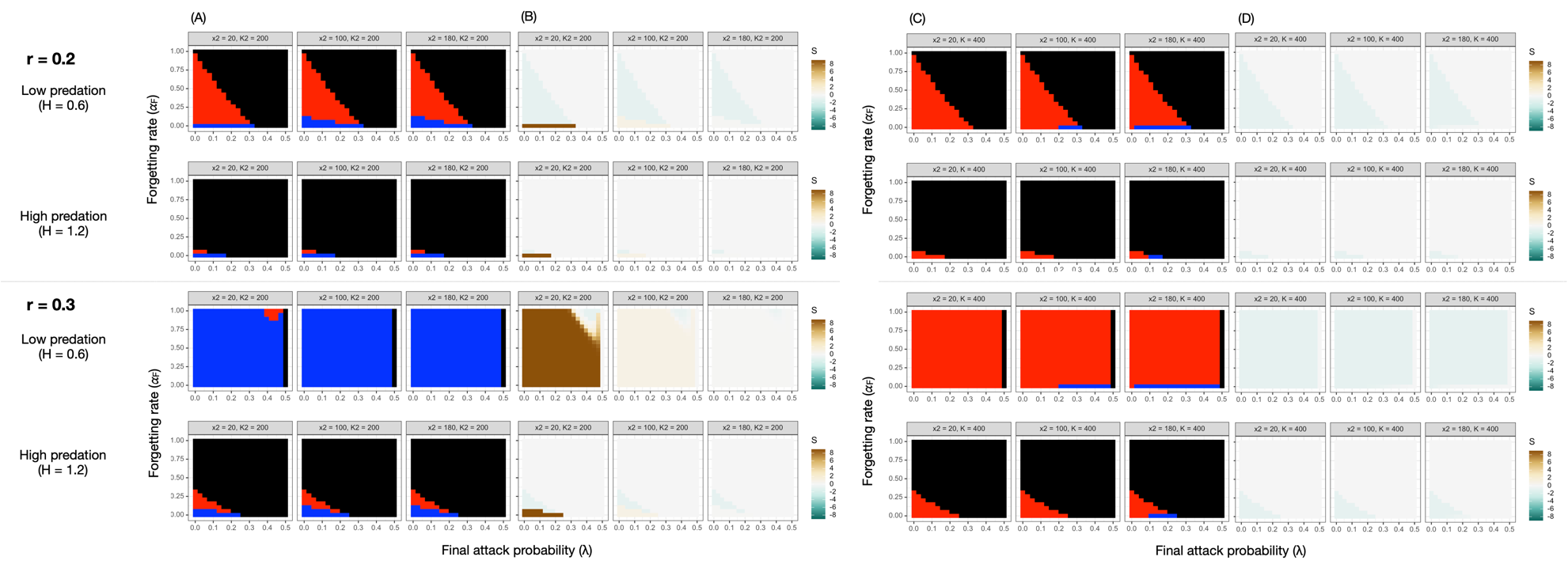

**Figure S2.** Signal persistence (A, C) and corresponding selection (B, D) on the rare signal, with respect to final attack probability (𝜆, x-axis) and forgetting rate (𝛼_F_, y-axis). Plots are organized based on the intrinsic rate of increase (*r*), per capita predation pressure (*H*). Columns in (A-D) are results when the rare signal was initially 10%, 50%, and 90% as abundant as the common signal. Black cells in (A) and (C) denote the extinction of both signals, red cells denote the extinction of only the rare signal, and blue cells denote the persistence of both signals. In (B) and (D), green colors indicate negative selection on the rare signal, brown colors indicate positive selection, and gray color denotes a selection value of zero. (A) and (B) are when prey have separate carrying capacities, and (C) and (D) are when they share the same carrying capacity. The results here are from simulations where the carrying capacity of the rare prey were set to be the same as the common prey despite its initial minority.

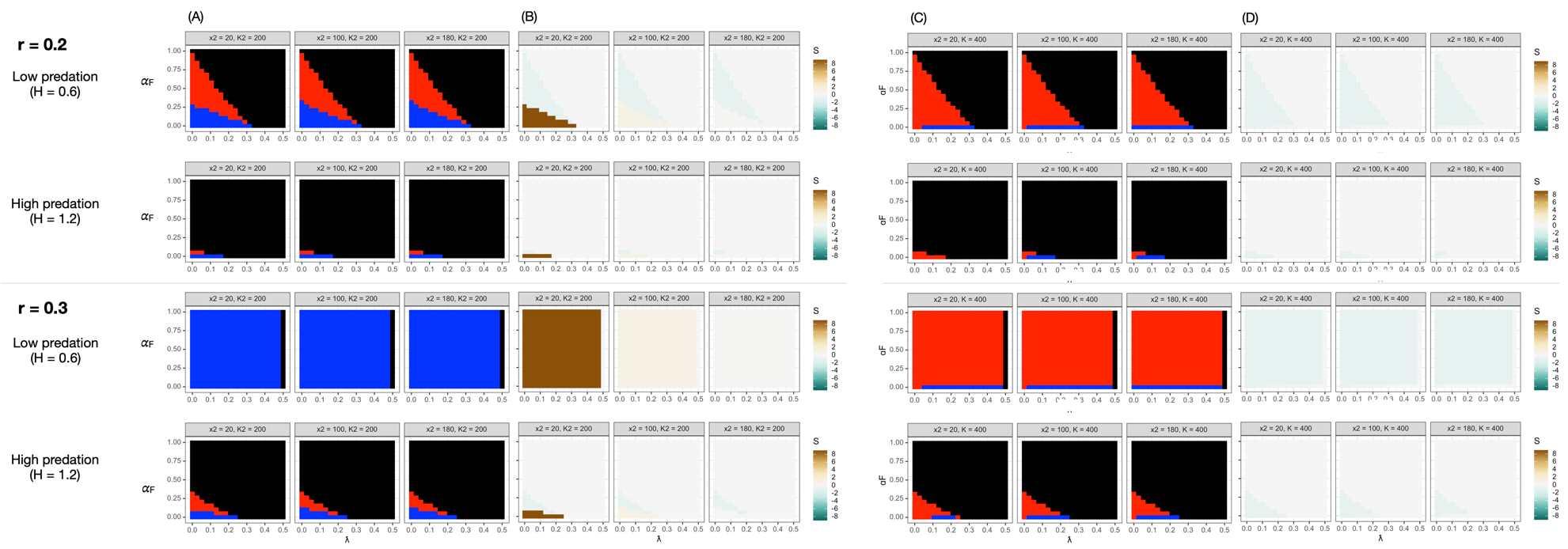

**Figure S3.** Signal persistence (A, C) and corresponding selection on the rare signal (B, D), with respect to intrinsic rate of increase (*r*), per capita predation pressure (*H*), and the initial abundance of the rare signal (10%, 50%, and 90% relative to the common signal) when there is generalization (G = 0.5, see main text). Generalization facilitated local warning signal diversity in all scenarios, especially when the ecological conditions were already favorable (e.g., high intrinsic rate of increase and low per capita predation pressure). The organization of the plots and coloring of the cells are the same as in Figure S2. . (A) and (B) are when prey have separate carrying capacities, and (C) and (D) are when they share the same carrying capacity. As in Figure S2, results are from simulations where the rare prey could be as abundant as the common prey despite their initial rarity.

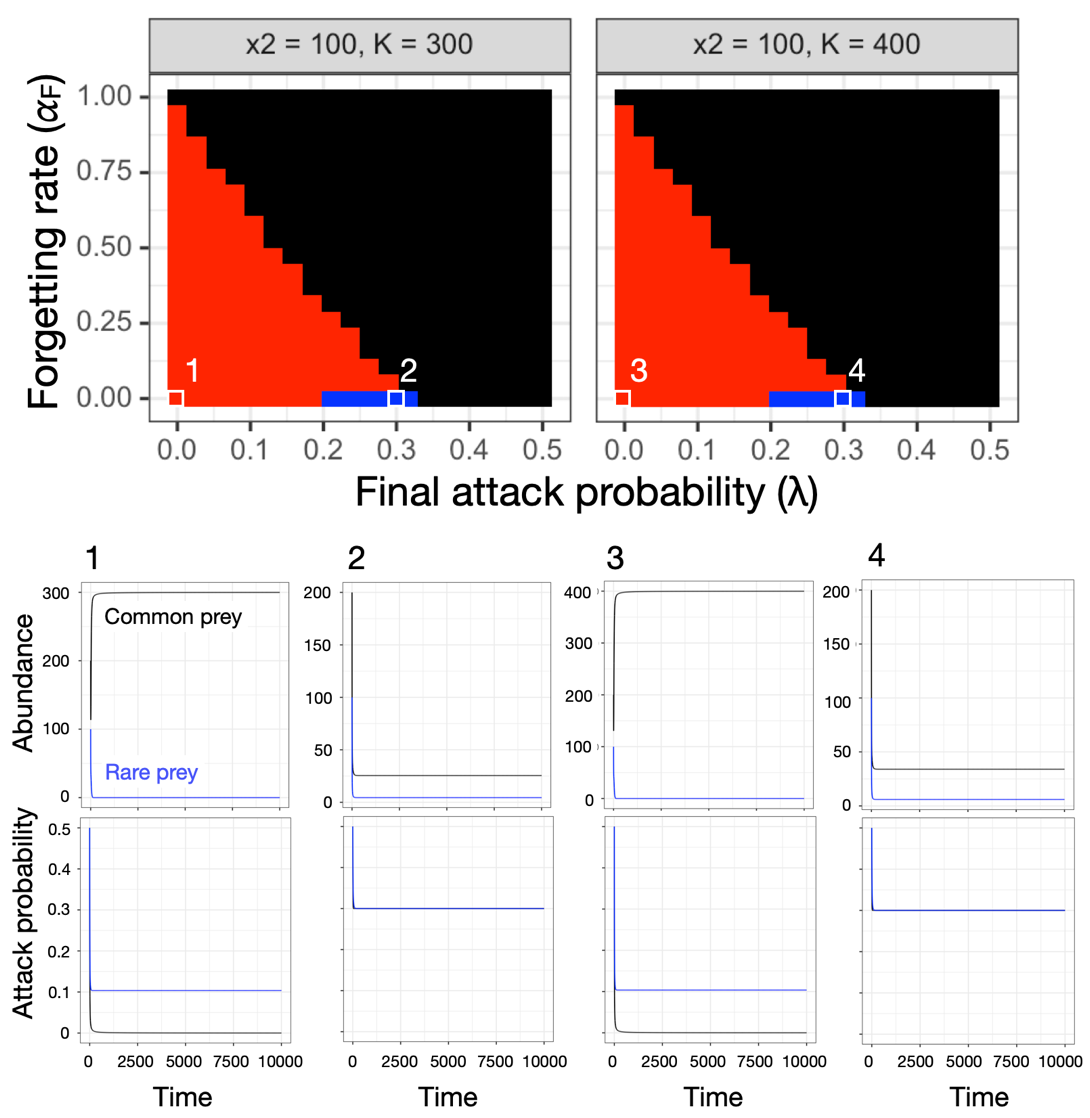

**Figure S4.** Signal persistence under a shared prey carrying capacity. Rare signal were driven to extinction in conditions 1 and 3, in which predation was the weakest due to high predator avoidance and long-lasting memory. In conditions 2 and 4, predation offset competitive disadvantage against the rare signal under low predator forgetting rate and intermediate values of final attack probability. The results were qualitatively identical with higher shared carrying capacities; only the steady-state prey abundances differed.

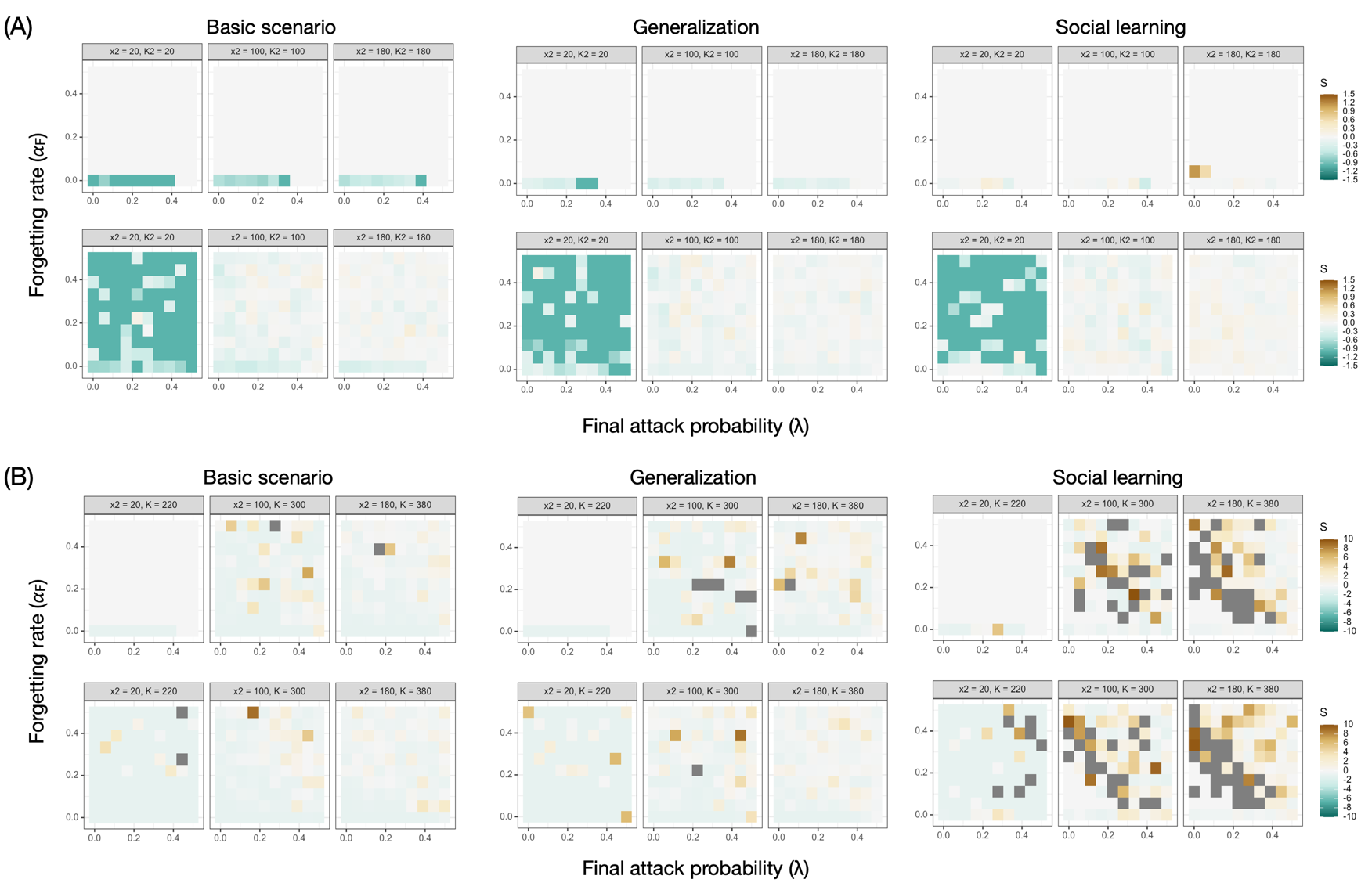

**Figure S5.** Selection on the rare signal with respect to final attack probability and forgetting rate in the individual-based simulations under the basic scenario, with generalization, and with social learning. (A) Selection on the rare signal when prey have separate carrying capacities (B) Selection on the rare signal when prey share the same carrying capacity. Within (A) and (B), plots in the top panel are from simulations with *r* = 0.2, and those in the bottom panel are with *r* = 0.3. Dark gray denotes selection values on the rare signal higher than 10.

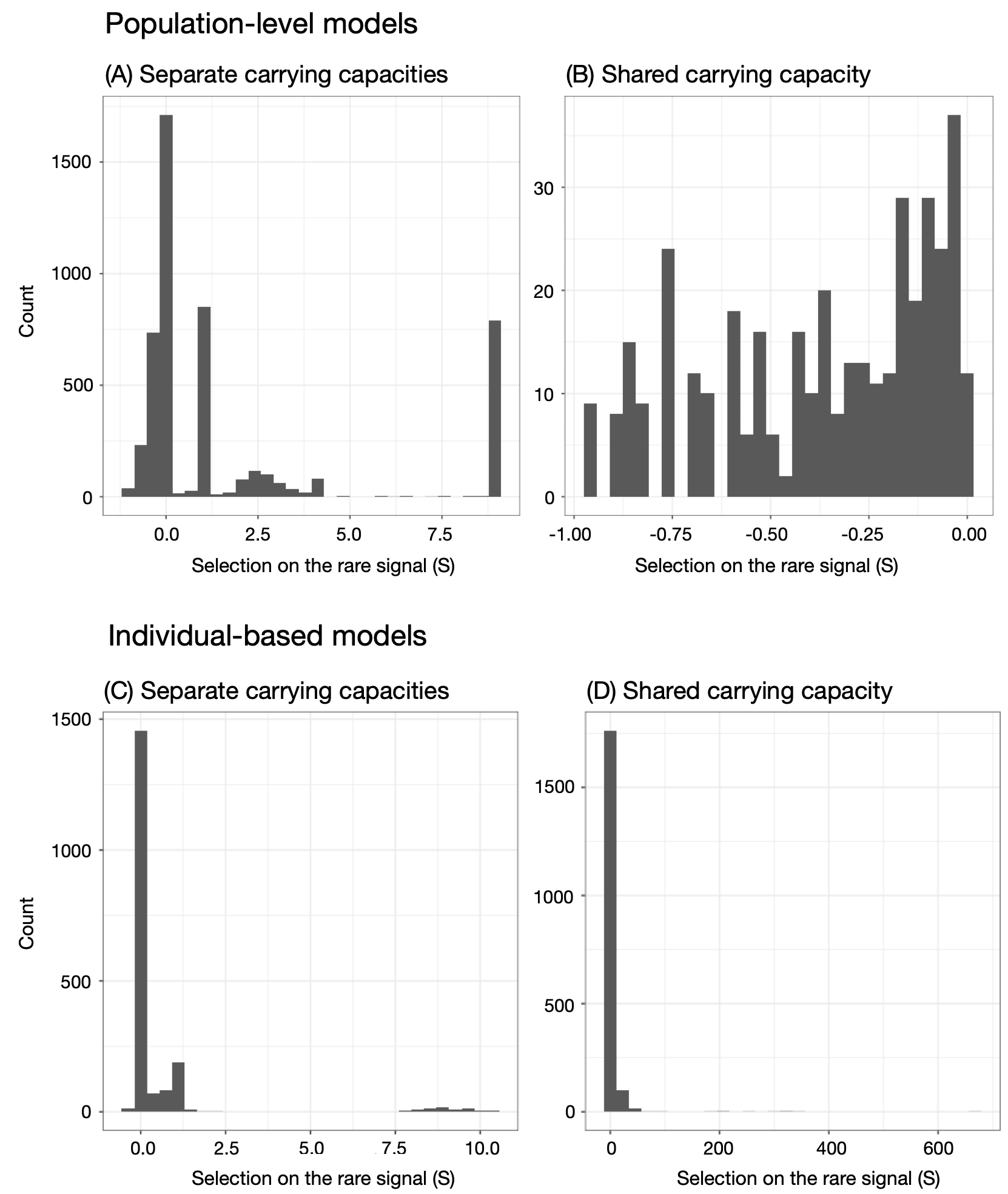

**Figure S6.** Distribution of selection values on the rare signal when it persisted in the population-level model (A, B) and the individual-based models (C, D), when prey had separate or shared carrying capacities.

**Table S1.** Empirically measured final attack probability ($\lambda$) and forgetting rate (𝛼_F_) from selected predators used in Figure 2.

| **Species** | **Taxon** | $\boldsymbol{\lambda}$ | $\alpha_{F}$ | **Source** |
| --- | --- | --- | --- | --- |
| *Habronattus pyrrithrix* | Arachnid | 0 | 0.011 | (Taylor *et al.* 2016) |
| *Thalassoma bifasciatum* | Fish | 0.05 | 0.5 | (Miller & Pawlik 2013) |
| *Thalassoma bifasciatum* | Fish | 0.165 | 0.5 | (Miller & Pawlik 2013) |
| *Galls gallus domesticus* | Bird | 0.256 | 0.006 | (Siddall & Marples 2008) |
| *Parus major* | Bird | 0.118 | 0.009 | (Lindström *et al.* 2006) |
| *Parus major* | Bird | 0.119 | 0.026 | (Lindström *et al.* 2006) |
| *Parus major* | Bird | 0.051 | 0.008 | (Lindström *et al.* 2006) |
| *Chalcides sexlineatus* | Lizard | 0.083 | 0.035 | (Beneš & Veselý 2017) |
| *Planigale maculate* | Mammal | 0 | 0.0003 | (Webb *et al.* 2008) |

**Table S2.** Strengths of generalization and social learning from published studies

| Source | Taxon | Species | Value |
| --- | --- | --- | --- |
| **Generalization** | | | |
| (Raška *et al.* 2020) | Arachnid | *Evarcha arcuate* | 0.547 |
| (Raška *et al.* 2020) | Arachnid | *Evarcha arcuate* | 0.708 |
| (Taylor *et al.* 2016) | Arachnid | *Habronattus pyrrhithrix* | 0 |
| (Paez *et al.* 2021) | Bird | *Cyanistes caeruleus* | 0 |
| (Paez *et al.* 2021) | Bird | *Cyanistes caeruleus* | 0 |
| (Paez *et al.* 2021) | Bird | *Cyanistes caeruleus* | 0.256 |
| (Paez *et al.* 2021) | Bird | *Cyanistes caeruleus* | 0.344 |
| (Paez *et al.* 2021) | Bird | *Cyanistes caeruleus* | 0.562 |
| (Paez *et al.* 2021) | Bird | *Cyanistes caeruleus* | 1 |
| (Rönkä *et al.* 2018) | Bird | *Cyanistes caeruleus* | 0 |
| (Aronsson & Gamberale-Stille 2008) | Bird | *Gallus gallus domesticus* | 0 |
| (Aronsson & Gamberale-Stille 2008) | Bird | *Gallus gallus domesticus* | 0 |
| (Aronsson & Gamberale-Stille 2008) | Bird | *Gallus gallus domesticus* | 1 |
| (Aronsson & Gamberale-Stille 2008) | Bird | *Gallus gallus domesticus* | 1 |
| (Aronsson & Gamberale-Stille 2009) | Bird | *Gallus gallus domesticus* | 0.910 |
| (Aronsson & Gamberale-Stille 2009) | Bird | *Gallus gallus domesticus* | 0.947 |
| (Aronsson & Gamberale-Stille 2009) | Bird | *Gallus gallus domesticus* | 0.970 |
| (Aronsson & Gamberale-Stille 2009) | Bird | *Gallus gallus domesticus* | 1 |
| (Ihalainen *et al.* 2008b) | Bird | *Parus major* | 0 |
| (Ihalainen *et al.* 2008b) | Bird | *Parus major* | 0 |
| (Ihalainen *et al.* 2008b) | Bird | *Parus major* | 0.285 |
| (Ihalainen *et al.* 2008a) | Bird | *Parus major* | 0 |
| (Ihalainen *et al.* 2008a) | Bird | *Parus major* | 0 |
| (Ihalainen *et al.* 2008a) | Bird | *Parus major* | 0 |
| (Ihalainen *et al.* 2008a) | Bird | *Parus major* | 0 |
| (Lindström *et al.* 2006) | Bird | *Parus major* | 0 |
| (Lindström *et al.* 2006) | Bird | *Parus major* | 0 |
| (Lindström *et al.* 2006) | Bird | *Parus major* | 0 |
| (Lindström *et al.* 2006) | Bird | *Parus major* | 0 |
| (Miller & Pawlik 2013) | Fish | *Thalassoma bifasciatum* | 0 |
| (Miller & Pawlik 2013) | Fish | *Thalassoma bifasciatum* | 1 |
| (Beneš & Veselý 2017) | Lizard | *Chaldes sexlineatus* | 0.444 |
| **Social learning** | | | |
| (Salva *et al.* 2009) | Bird | *Gallus gallus domesticus* | 0.304 |
| (Hämäläinen *et al.* 2019) | Bird | *Parus major* | 0.306 |
| (Hämäläinen *et al.* 2020) | Bird | *Parus major* | 0.351 |
| (Hämäläinen *et al.* 2020) | Bird | *Parus major* | 0.416 |
| (Hämäläinen *et al.* 2020) | Bird | *Cyanistes caeruleus* | 0.551 |
| (Hämäläinen *et al.* 2020) | Bird | *Cyanistes caeruleus* | 0.689 |
